# The EGFR/ErbB inhibitor neratinib modifies the neutrophil phosphoproteome and promotes apoptosis and clearance by airway macrophages

**DOI:** 10.1101/2022.04.14.488364

**Authors:** Kimberly D Herman, Carl G Wright, Helen M Marriott, Sam C McCaughran, Mark O Collins, Stephen A Renshaw, Lynne R Prince

**Affiliations:** Department of Infection, Immunity and Cardiovascular Disease, The Medical School, Beech Hill Road, Sheffield, S10 2RX; The Bateson Centre, University of Sheffield, Western Bank, Sheffield, S10 2TN; Department of Biomedical Science, University of Sheffield, Sheffield, S10 2TN, UK

## Abstract

Dysregulated neutrophilic inflammation can be highly destructive in chronic inflammatory diseases due to prolonged neutrophil lifespan and continual release of histotoxic mediators in inflamed tissues. Therapeutic induction of neutrophil apoptosis, an immunologically silent form of cell death, may be beneficial in these diseases, provided that the apoptotic neutrophils are efficiently cleared from the tissue. Our previous research identified ErbB inhibitors as able to induce neutrophil apoptosis and reduce neutrophilic inflammation both *in vitro* and *in vivo* (Rahman et al., 2019). Here we extend that work using a clinical ErbB inhibitor, neratinib, which has the potential to be repurposed in inflammatory diseases. We show that neratinib reduces neutrophilic migration to an inflammatory site in zebrafish larvae. Neratinib upregulates efferocytosis and reduces the number of persisting neutrophil corpses in mouse models of acute, but not chronic, lung injury, suggesting the drug may have therapeutic benefits in acute inflammatory settings. Phosphoproteomics analysis of human neutrophils shows that neratinib modifies the phosphorylation of proteins regulating apoptosis, migration and efferocytosis. This work identifies a potential mechanism for neratinib in treating acute lung inflammation by upregulating the clearance of dead neutrophils and, through examination of the neutrophil phosphoproteome, provides important insights into the mechanisms by which this may be occurring.

## Introduction

Neutrophils are innate immune cells, crucial for protecting against infectious insults, and are a key cellular driver of the acute inflammatory response. If acute inflammation does not resolve however, the continual release of inflammatory mediators such as proteases and oxidative molecules by neutrophils can be highly histotoxic and prevent tissue healing, resulting in chronic inflammation. Neutrophils are considered a major contributor to irreversible lung damage in chronic obstructive pulmonary disease (COPD), in which chronic inflammation in the lungs is induced by continual exposure to noxious substances such as cigarette smoke and pollution (Barnes, 2017). Current treatments for COPD focus on reducing symptom severity and none alter disease progression (Vogelmeier et al., 2017). Targeting the underlying neutrophilic inflammation may bring much needed new approaches for the third leading cause of death worldwide (World Health Organisation, 2019).

Neutrophils are one of the shortest-lived cells in the body and typically die by spontaneous apoptosis, an immunologically silent cell death mechanism in which intracellular proteins are degraded to prevent further function, but the cell membrane remains intact (Fox et al., 2010). Apoptotic neutrophils are rapidly ingested by phagocytes such as macrophages in a process called efferocytosis. If this does not occur efficiently, apoptotic neutrophils can undergo secondary necrosis, in which the cell membrane ruptures and the highly inflammatory intracellular contents spills onto the tissue, inducing further inflammation (Szondy et al., 2017). Neutrophil apoptosis is known to be delayed in inflammatory environments, including in the lungs of patients with COPD during exacerbations (acute worsening of symptoms), resulting in the further release of histotoxic contents and tissue damage (Brown et al., 2009; Pletz et al., 2004). Upregulating neutrophil apoptosis may therefore be beneficial in chronic inflammatory diseases to prevent such damage, however it would be important that these dead cells are efficiently removed from the tissue.

Previous research by our group showed that the inhibitors of the epidermal growth factor receptor (EGFR) or ErbB family of receptor tyrosine kinases are able to accelerate neutrophil apoptosis and reduce neutrophilic inflammation in several experimental models (Rahman et al., 2019). Here we have built on that work by investigating the efficacy of neratinib, a clinical ErbB inhibitor used to treat breast cancer, in mouse models of acute and chronic lung inflammation. Neratinib is known to be safe and tolerated by humans (Chan et al., 2021), making it an attractive therapeutic for repurposing.

We have also investigated the mechanism by which neratinib induces apoptosis of human neutrophils. In normal development, ErbB signalling regulates pathways controlling cellular transcription, proliferation, survival, migration and differentiation, whereas excessive ErbB signalling in some tumour cells inhibits normal apoptosis and promotes oncogenesis (Roskoski, 2014). Pharmacological ErbB inhibitors induce tumour cell apoptosis by blocking these aberrantly activated signalling pathways; however ErbBs are not known regulators of neutrophil apoptosis, and their role in neutrophils and other immune cells is sparsely studied. ErbBs are kinases and the regulation of their downstream signalling pathways, like many intracellular signalling networks, is controlled in part by protein phosphorylation. We therefore interrogated the “phosphoproteome” of neutrophils to obtain insights into the mechanisms by which neratinib is exerting its effects.

## Results

### Neratinib treatment reduces neutrophilic inflammation in a larval zebrafish injury model, and induces apoptosis of human neutrophils *in vitro*

Research in our previous paper used a larval zebrafish model of injury-induced inflammation (Renshaw et al., 2006) to test the effect of ErbB inhibitors on neutrophils in an inflammatory environment. We showed that treatment of larvae with the research grade ErbB inhibitors tyrphostin AG825 and CP-724,714 reduced neutrophil number at the tail fin injury site (Rahman et al., 2019). Here we tested the clinical ErbB inhibitor, neratinib, to determine if it is similarly effective at reducing neutrophilic inflammation in this model. Using the transgenic zebrafish neutrophil reporter line *TgBAC(mpx:EGFP)i114* (Figure 1A), we found that 16 hour pre-treatment of larvae with 10 μM neratinib reduced the number of neutrophils at the injury site at both 4 and 8 hours post injury (Figure 1B). We also found that the total number of neutrophils across the whole body of zebrafish larvae was unchanged with neratinib treatment, suggesting that neratinib is acting specifically on the neutrophilic response to inflammation (Figure 1C).

**Figure 1.**
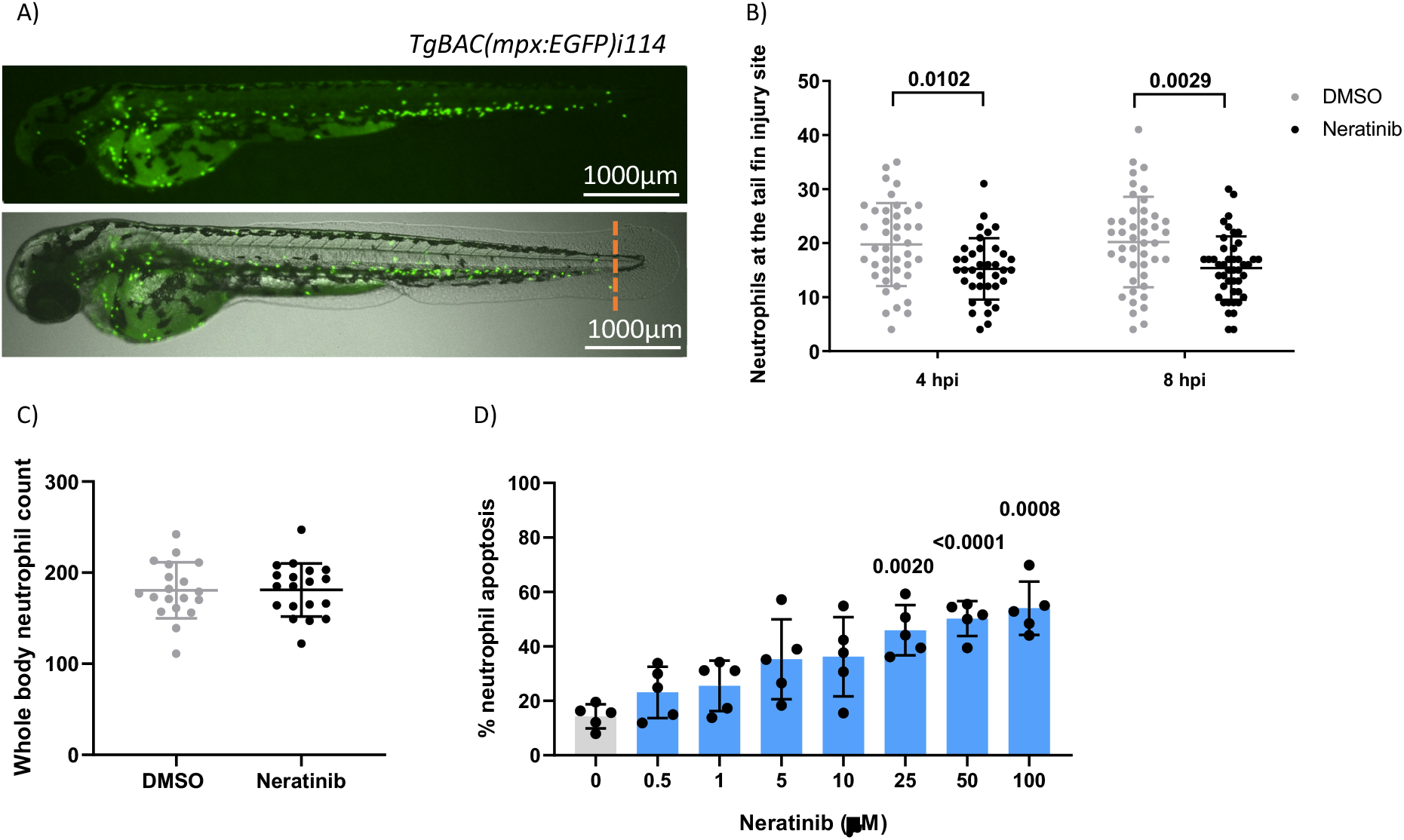
Neratinib reduces neutrophilic inflammation at the tail fin injury site of zebrafish larvae in vivo, and induces apoptosis of human neutrophils in vitro. Neutrophils in zebrafish larvae can be enumerated by fluorescence microscopy using the transgenic line *TgBAC(mpx:EGFP)i114*, in which each GFP-expressing (green) cell is counted as one neutrophil (A). Tail fin transection (A, orange dotted line) was performed after 16h treatment with neratinib, and neutrophils at the site of injury enumerated 4- and 8-hours post-injury. Larvae treated with neratinib had reduced numbers of neutrophils at the tail fin injury site at both timepoints in comparison to DMSO-treated larvae (B). Minimum n=30 larvae per condition across three experimental repeats, data analysed by two-way ANOVA with Sidak’s multiple comparisons. Total neutrophil number across the whole body of larvae was unchanged after 16h treatment with 10 μM neratinib, in comparison to control DMSO-treated larvae (C). N=20 larvae per condition across three independent experiments, data analysed by unpaired t test. In human neutrophils isolated from whole blood samples from healthy volunteers, treatment with neratinib *in vitro* results in a dose-dependent increase in the rate of apoptosis (D). N=5 healthy blood donors, data analysed by one-way ANOVA with Dunnett’s multiple comparisons, comparing each neratinib concentration with the DMSO control treatment. Each data point represents data from one larva or human blood donor. Bars show mean ± standard deviation. P values indicated where p<0.05.

We have previously shown that a number of ErbB inhibitors accelerate the rate of apoptosis of human neutrophils *in vitro*, and here we confirm this using neratinib. Neratinib treatment was shown to induce apoptosis after 6 hours of treatment, reaching significance at 25μM (Figure 1D). These findings show that this clinical ErbB inhibitor also promotes human neutrophil apoptosis *in vitro*, at similar concentration ranges as research-grade ErbB inhibitors.

### Phosphoproteomics analysis of human neutrophils reveals potential mechanisms of action for neratinib

Although ErbB signalling has been studied widely in fields such as development and cancer, details of these signalling pathways in neutrophils, or any other immune cells, is sparse in the literature. Since neutrophils are one of the shortest-lived cells in the human body and rapidly undergo spontaneous apoptosis, we considered that the mechanism by which neratinib induces neutrophil apoptosis may be different to that described in other cell types. As protein phosphorylation is the key event downstream of receptor tyrosine kinases and regulates many cellular processes, we investigated this using mass spectrometry-based differential phosphoproteomics to identify changes in the neutrophil phosphoproteome that occur with neratinib treatment.

For this experiment, neutrophils were isolated from the blood of five healthy volunteers and treated with either 25μM neratinib or equivalent volume of DMSO (control) for 1 hour, followed by a cell-permeable analogue of cyclic AMP; dibutyryl-cAMP (db-cAMP) for 30 minutes. Cyclic AMP supresses apoptosis in human neutrophils (Martin et al., 2001; Rahman et al., 2019), switching on pro-survival signalling pathways that, based on previous findings, we hypothesise will be supressed in neratinib treated cells (Rahman et al., 2019). Neutrophil proteins were isolated and digested, and enriched phosphopeptides were identified and quantified using LC-MS/MS analysis. A single DMSO-treated sample was excluded from analysis due to poor correlation with other samples.

Details of all phosphopeptides detected, including information about specific phosphorylation events, are detailed in Supplementary File 1. Since all samples were treated with db-cAMP, it was expected that phosphorylated peptides mapping to proteins downstream of cAMP signalling pathways would be detected. Protein kinase A is directly activated by cAMP (Sassone-Corsi, 2012), and several phosphopeptides mapping to subunits of this protein complex were identified in the dataset (Figure 2 - figure supplement 1). Other phosphopeptides mapping to downstream proteins present in the dataset included BRAF, CREB, GSK3A, and several MAPK family members. The detection of these phosphorylated peptides in the neutrophil samples was considered an effective validation of the dataset.

To determine if neratinib was modifying the phosphorylation of proteins within the ErbB signalling pathways described in literature, we interrogated the dataset for downstream components of ErbB signalling and as above. A key downstream mediator of ErbB signalling is the PI3K family. Phosphopeptides mapping to several different subunits of this family were identified (PIK3AP1, PIK3R1, PIK3R5), which were detected in similar numbers of samples in both treatment groups (Figure 2 - figure supplement 2). Similar results were observed for members of the AKT family and several members of the MAPK family. Although there are differences in the detection of some phosphopeptides between the treatment groups, such as STAT3 which was detected in 2/5 neratinib treated samples and all DMSO treated samples, the majority of phosphorylated proteins downstream of ErbB signalling were not differentially detected (Figure 2 - figure supplement 2), suggesting neratinib is not modifying ErbB signalling pathways that are described in literature.

We then used an unbiased approach to identify differences in the phosphoproteome of neratinib vs DMSO treated neutrophils. Statistical analysis was used to determine if the abundance of any phosphopeptide was different between the two treatment groups. At 5% false discovery rate, 16 phosphorylated peptides mapping to 15 proteins were identified as statistically regulated (Supplementary File 1, statistical analysis tabs), with 9 phosphopeptides mapping to 8 proteins statistically increased in the DMSO treatment group and 7 in the neratinib treatment group (Figure 2 - figure supplement 3A). This dataset of 15 proteins was input into the online tool STRING, which identifies interactions between proteins. One interaction was identified between MAML1 and NCOR1, which were both more abundant in neratinib treated samples (Figure 2 - figure supplement 3B), however no other proteins in this dataset were found to interact.

In order to define a larger set of phosphorylation sites putatively regulated by neratinib, we combined sites identified by statistical analysis with those that were enriched in either treatment group i.e. phosphorylation sites identified in DMSO treated samples but not neratinib samples, and vice versa. All phosphopeptides detected in 3-4 DMSO and 0-1 neratinib treated samples were considered enriched with DMSO treatment, and all phosphopeptides detected in 4-5 neratinib and 0-1 DMSO-treated samples were considered enriched with neratinib treatment. This equated to 70 DMSO-enriched and 95 neratinib-enriched phosphorylated peptides, listed in Supplementary File 1 (tabs 2 and 3, respectively), which mapped to 62 DMSO-enriched and 81 neratinib-enriched phosphorylated proteins. These two datasets, combined with the phosphopeptides that were identified as statistically more abundant in either treatment group, were analysed using STRING (Figure 2A-B).

**Figure 2.**
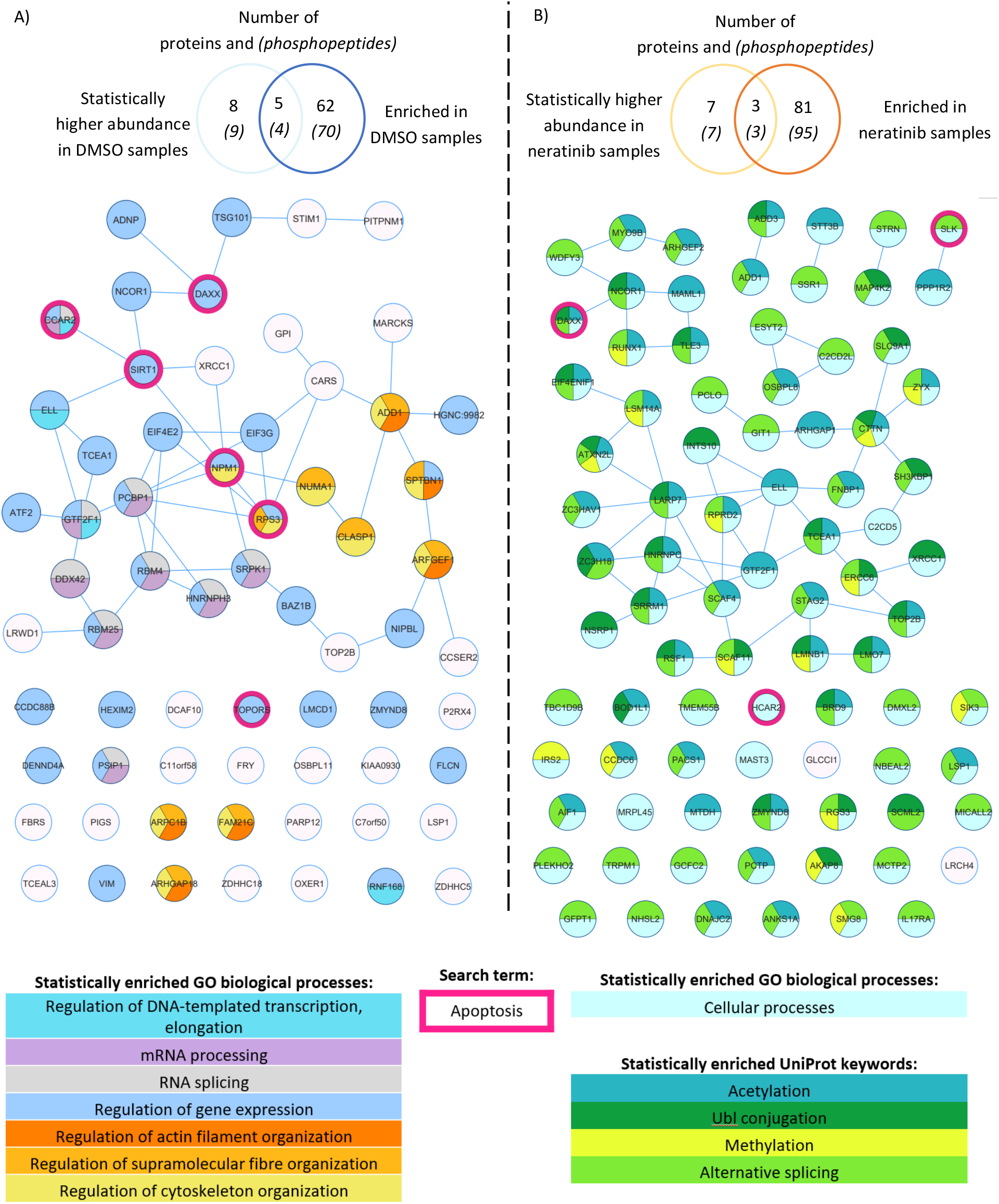
Phosphoproteomics analysis of human neutrophils show that neratinib treatment induces changes in phosphorylated proteins regulating numerous biological processes. STRING identified interactions between phosphorylated proteins in the combined “DMSO-enriched” and “statistically higher abundance in DMSO” datasets (A), and the combined “neratinib-enriched” and “statistically higher abundance in neratinib” datasets (B). Venn diagrams show the number of phosphopeptides and proteins they map to in each dataset, and the overlapping proteins between the two datasets for each treatment group. STRING analysis of these datasets indicates interactions between proteins (lines). STRING also identified a number of Gene Ontology (GO) biological processes that were statistically enriched in both treatment groups, highlighted in colour. The neratinib-enriched treatment group had only one statistically enriched biological process, and so statistically enriched keywords from the UniProt database are also shown. Both datasets were searched for the keyword “apoptosis”, and hit proteins highlighted with a pink outline.

STRING identified several functionally enriched biological processes (extracted from Gene Ontology), i.e. processes that are observed within these networks more frequently than expected based on hypergeometric testing (Gene Ontology Consortium, 2021; Szklarczyk et al., 2021) (Figure 2). In DMSO-treated samples, a number of enrichments related to gene expression, e.g. mRNA processing and regulation of DNA-templated transcription and elongation, as well as regulation of actin filament, supramolecular fibre and cytoskeleton organisation (Figure 2A). These proteins might not be phosphorylated in neratinib-treated cells due to the onset of apoptosis and subsequent shutting down of these cellular processes. Within neratinib-treated samples, the only enriched biological process identified by STRING was “cellular processes” (Figure 2B). Several keywords extracted from the protein database UniProt were also identified as statistically enriched, including methylation, alternative splicing, and ubiquitin-like protein (Ubl) conjugation (Figure 2B). The latter is a key step in proteolysis, and these proteins are possibly phosphorylated in neratinib-treated cells due to the onset of apoptosis. In addition to analysing the DMSO- and neratinib-treated samples separately, the datasets were combined and entered into STRING for analysis, as a phosphorylation of one protein may result in the removal of a phosphate group from another (for example phosphatases that are regulated by phosphorylation). Biological processes enriched in this combined dataset were similar to that in the DMSO-treated samples, but also included regulation of protein polymerisation and mRNA metabolic processes (Figure 2 - figure supplement 4). Analysis of this dataset using the online tool Reactome (Fabregat et al., 2017) identified the top significant enrichment as being the Rho GTPase cycle, which in neutrophils and other cell types regulate migration, phagocytosis and efferocytosis by controlling cytoskeletal arrangements (Kim et al., 2017; McCormick et al., 2019).

Apoptosis was not identified as a statistically enriched process or keyword in any of the datasets analysed, however several proteins within both datasets have apoptosis listed as a UniProt keyword. These include NPM1, TOPORS and SIRT1 which regulate p53 activity, and RBM25 which regulates the ratio of pro-and anti-apoptotic BCL2 isoforms (Figure 2, pink circles). These may be driving the pro-apoptotic mechanism of neratinib in human neutrophils. Two of these candidates, NPM1 and SIRT1, were tested to determine whether they induce apoptosis via the same mechanism as neratinib in human neutrophils, by assessing the ability of pharmacological inhibitors to induce neutrophil apoptosis, alone or in combination with neratinib. A pharmacological inhibitor of SIRT1 did not induce neutrophil apoptosis (Figure 2 - figure supplement 5A), which perhaps suggests it is unlikely to be a candidate. Inhibiting NPM1 however, significantly increased neutrophil apoptosis (Figure 2 - figure supplement 5B). The combination of the NPM1 inhibitor and neratinib did not result in an additional increase in apoptosis in comparison to the NPM1 inhibitor alone, suggesting an epistatic relationship.

### Neratinib treatment increases macrophage efferocytosis and reduces numbers of neutrophil corpses in a murine model of LPS-induced acute lung injury

To determine whether neratinib may ultimately be beneficial as a treatment for patients with chronic inflammatory diseases such as COPD, we used several murine lung injury models. Our previous research showed that the tyrphostin AG825 increased neutrophil apoptosis and efferocytosis by macrophages in an acute LPS-induced lung injury model. We used the same model to treat mice with either neratinib (20mg/kg) or vehicle (Figure 3A). After culling all mice 48 hours post-treatment, cells in bronchoalveolar lavage fluid (BAL) were analysed and identified as neutrophils, macrophages or lymphocytes based on morphology (Figure 3B). The total cell number in BAL from each mouse was unchanged with neratinib treatment (Figure 3C), as was the percentage of neutrophils, macrophages and lymphocytes (Figure 3D-F). Macrophage efferocytosis was quantified by identifying inclusions of dead cells or cell debris within macrophage vesicles (Figure 3G). As macrophages may have multiple vesicles containing inclusions, the total number of inclusions per 100 macrophages was calculated, as well as the percentage of macrophages containing any inclusions; both measures of efferocytosis were significantly increased with neratinib treatment (Figure 3H-I).

**Figure 3.**
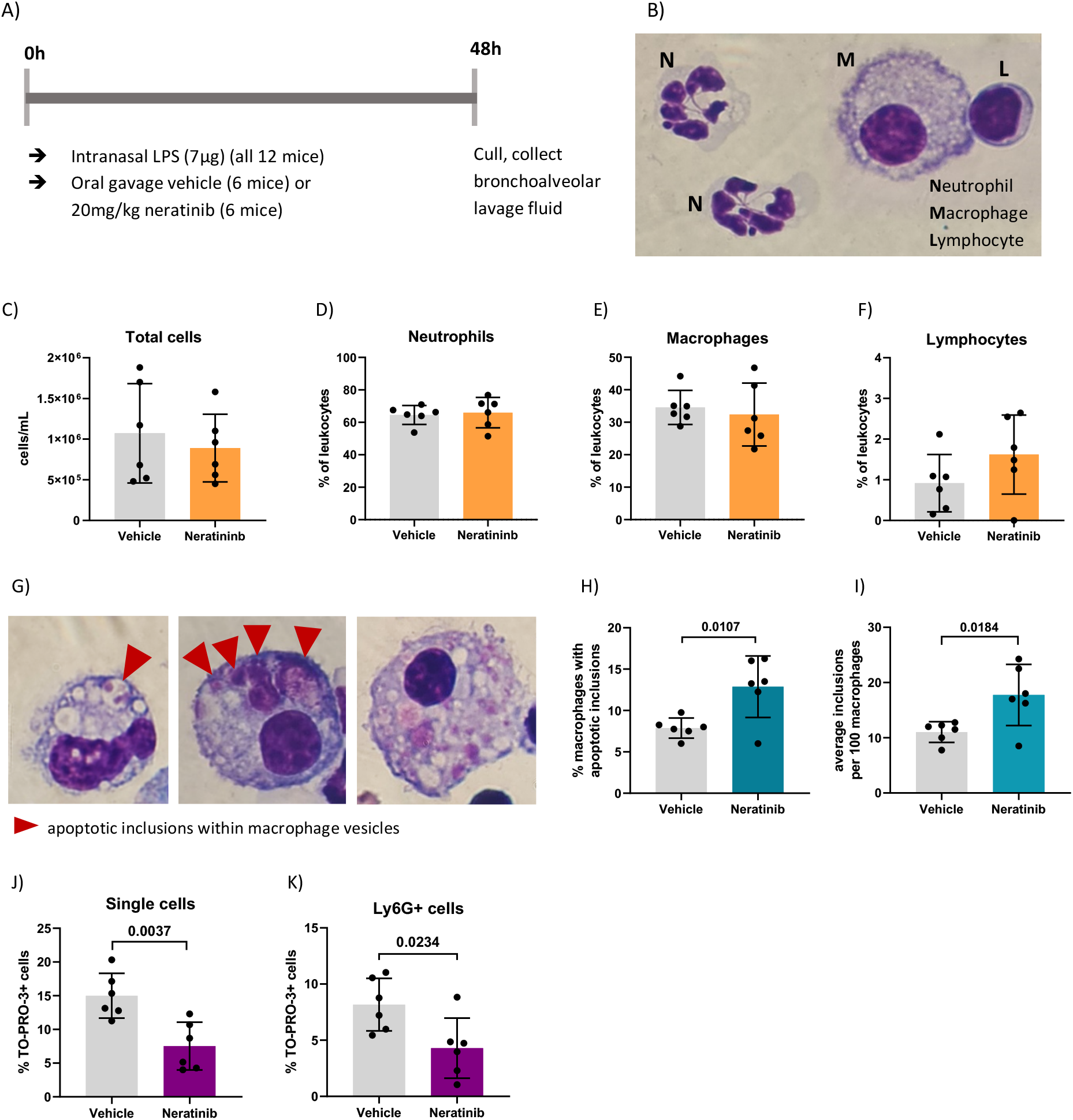
Neratinib treatment in a mouse LPS-induced acute lung injury model increases rates of macrophage efferocytosis, and reduces the number of neutrophil corpses in bronchoalveolar lavage fluid. Schematic of the treatment protocol (A). Cytospins of BAL were stained with Kwik-Diff and examined by light microscopy, and neutrophils, macrophages and lymphocytes identified by morphology (B). Total cells in BAL were counted using haemocytometer counting chamber (C), and the percentage of neutrophils (D), macrophages (E) and lymphocytes (F) in BAL were unchanged between treatment groups. The engulfment of cell debris by macrophages is visible as inclusions within intracellular vesicles, which can be identified with Kwik-Diff staining and light microscopy (G, red arrows). Cell debris may be broken into small pieces within a vesicle (G, left panel) or fill the vesicles (G, middle panel). Macrophages may contain one or more vesicle with inclusions, in some cases too many to count accurately (G, right panel); if above 6, the number of inclusions was recorded as 6. Both the percentage of macrophages containing any inclusions (H) and the total number of inclusions per 100 macrophages (I) was increased in the neratinib treatment group. Cells in BAL were also analysed by flow cytometry (gating strategy in Figure Supplement 1), and the percentage of TO-PRO-3+ cells (J) and TO-PRO-3+ neutrophils (Ly6G+ cells) (K), was reduced in neratinib treated mice. Each data point in graphs represents data from one mouse (n=6 per treatment group); bars show mean ± standard deviation. Unpaired t tests used for statistical analysis, p values indicated where p<0.05.

As we observed an increase in efferocytosis, we examined whether the percentage of free dead cells in BAL was altered with neratinib treatment. Cells were analysed by flow cytometry, staining for Ly6G to identify neutrophils, Annexin-V to identify apoptotic cells, and the viability dye, TO-PRO-3 (Figure 3 - figure supplement 1A). Since TO-PRO-3 can only enter cells that have lost membrane integrity (dead cells), these are referred to as cell corpses. The majority of TO-PRO-3+ cells also stained for Annexin-V (Figure 3 - figure supplement 1B-C), suggesting these cells either underwent apoptosis and subsequent secondary necrosis, or have directly lost membrane integrity. The percentage of both TO-PRO-3+ cells, and TO-PRO-3+ neutrophils was significantly decreased in the neratinib treatment group, supporting a role for neratinib enhanced efferocytosis in preventing the accumulation of cell corpses, and particularly neutrophils corpses (Figure 3J-K). Neratinib treatment did not alter the percentage of viable or apoptotic cells or neutrophils (Figure 3 - figure supplement 1D-G).

An increase in the rate of efferocytosis, and a decrease in the number of cell corpses, particularly neutrophil corpses, could be beneficial in the resolution of inflammation. We followed up these promising results in a chronic murine model of lung disease, to determine if neratinib is beneficial in a chronic inflammatory environment.

### Neratinib treatment reduces inflammatory cytokines in chronic murine lung injury model

To determine if neratinib may have therapeutic benefit in a chronic model of lung inflammation, mice were given weekly intranasal doses of LPS and elastase over 4 weeks, which is known to result in COPD-like features within 4 weeks (Ganesan et al., 2012). Elastase induces widespread tissue damage and is a profound inducer of emphysema-like damage and bronchitis. LPS is an excellent inflammatory stimulus and also models the contribution of infection (compared to isolated cigarette smoke models for example). Overall, these treatments induce airway inflammation, emphysematous changes and functional impairment comparable to COPD. We tested two different dosing schedules for neratinib administration: one in which neratinib was given at the same time as the LPS/elastase (induction dosing) (Figure 4A), and the second in which the neratinib treatment began 2 weeks after disease onset (therapeutic dosing) (Figure 4B). In both cases, mice were culled on day 28, and BAL and blood collected for analysis. All mice developed lung inflammation, with evidence of epithelial damage and areas of alveolar enlargement, as determined by histological analysis (data not shown).

**Figure 4.**
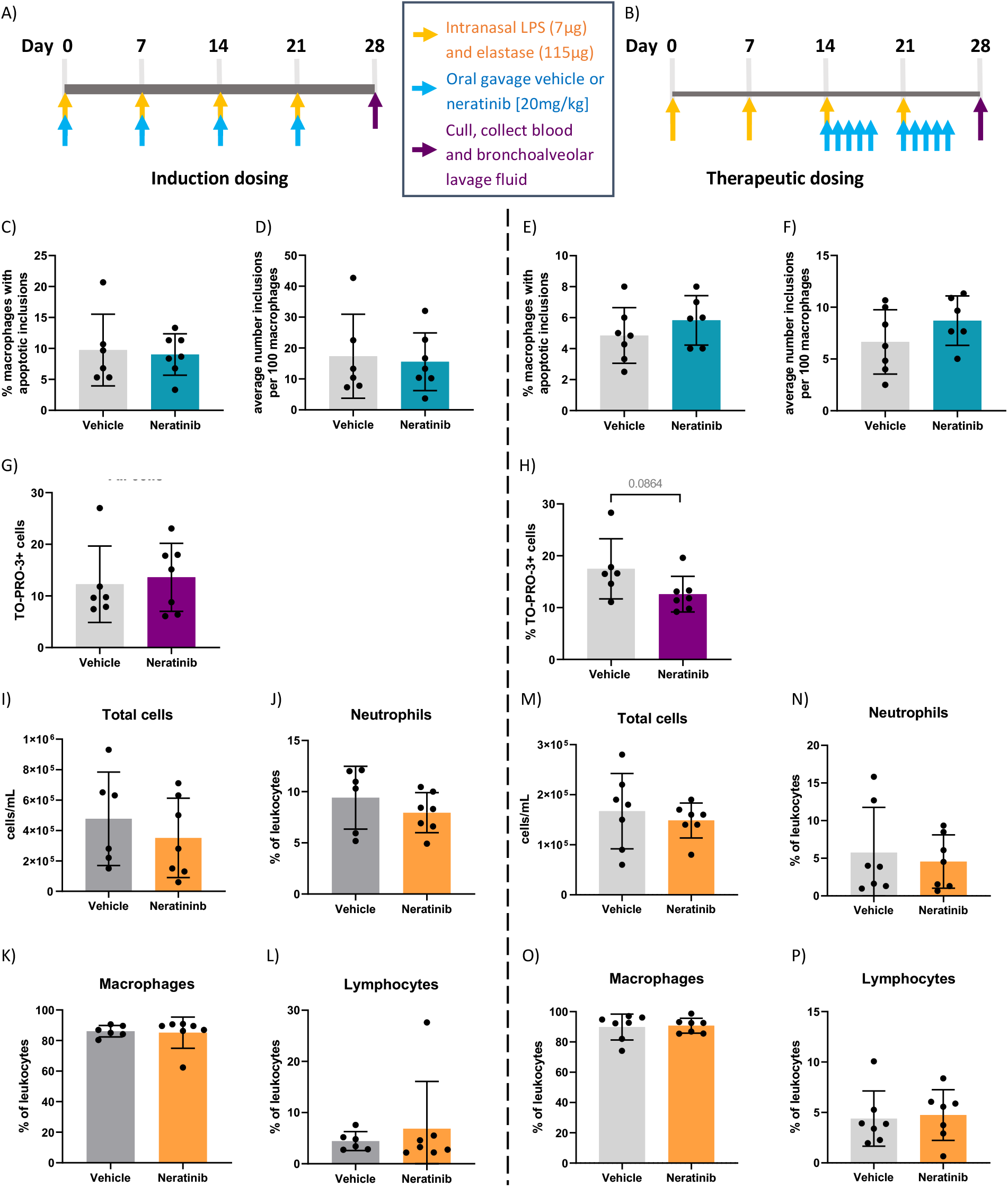
No changes in efferocytosis were observed with neratinib treatment in chronic models of lung disease, although dead cell numbers are reduced with a therapeutic dosing model. Two different dosing protocols were used for neratinib treatment in chronic lung disease model, induction dosing (A) and therapeutic dosing (B). Uptake of apoptotic cell debris by macrophages was unchanged between treatment groups in both the induction dosing (C, D) and therapeutic dosing (E, F) models. The percentage of TO-PRO-3+ cells in BAL, assessed by flow cytometry, was unchanged in the induction dosing model (G) and the therapeutic dosing model (H). Leukocyte number in BAL, and the percentage of neutrophils, macrophages and lymphocytes, were similarly unchanged across both models (I-P). Each data point in graphs represents data from one mouse; bars show mean ± standard deviation. Induction dosing study: n=6 vehicle treated mice, n=7 neratinib treated mice; therapeutic dosing study: n=7 mice in each treatment group. Unpaired t tests used for statistical analysis.

In contrast with the acute lung injury study, efferocytosis by macrophages was not significantly modified by neratinib treatment (Figure 4C-F). Cells within BAL were also analysed by flow cytometry, and neither neratinib treatment schedule altered the percentage of TO-PRO-3+ cells, although a trend for decreased cell corpses was observed in the therapeutic model (Figure 4G-H). The total number of leukocytes in BAL samples, and the percentage of neutrophils, macrophages and lymphocytes were similarly unchanged between the treatment groups in both studies (Figure 4I-P).

We analysed the concentration of cytokines in supernatant from BAL, to determine if neratinib might be reducing inflammatory cytokine levels. Both IL-6 and KC (also known as CXCL1 or GROα) were significantly reduced with the induction dosing method (Figure 5A-B), but not therapeutic dosing of neratinib (Figure 5C-D).

**Figure 5.**
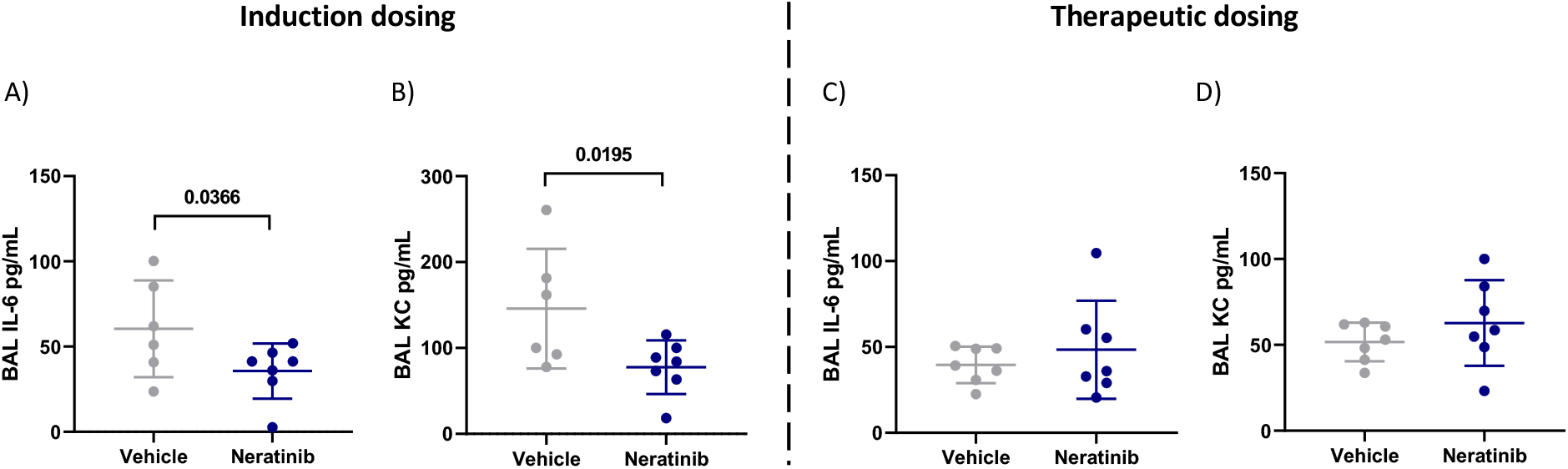
Neratinib treatment reduces levels of cytokines IL-6 and KC in BAL in the induction dosing model of lung disease. Cytokines were measured in BAL supernatant by ELISA. Interleukin-6 (IL-6) and KC were reduced in neratinib treated mice in the induction dosing model (A, B), whereas these were unchanged in the therapeutic dosing model (C, D). Each data point represents data from one mouse; bars show mean ± standard deviation. Unpaired t tests used for statistical analysis, p values indicated.

Blood samples were analysed using an automated haematology analyser, to enumerate circulating leukocyte numbers and differentiating between neutrophils, monocytes and lymphocytes. We found no changes in either the number of circulating leukocytes (Figure 6A-B) or in the percentage of neutrophils, monocytes and lymphocytes (Figure 6C-D) between the vehicle and neratinib treatment groups, with either dosing protocol.

**Figure 6.**
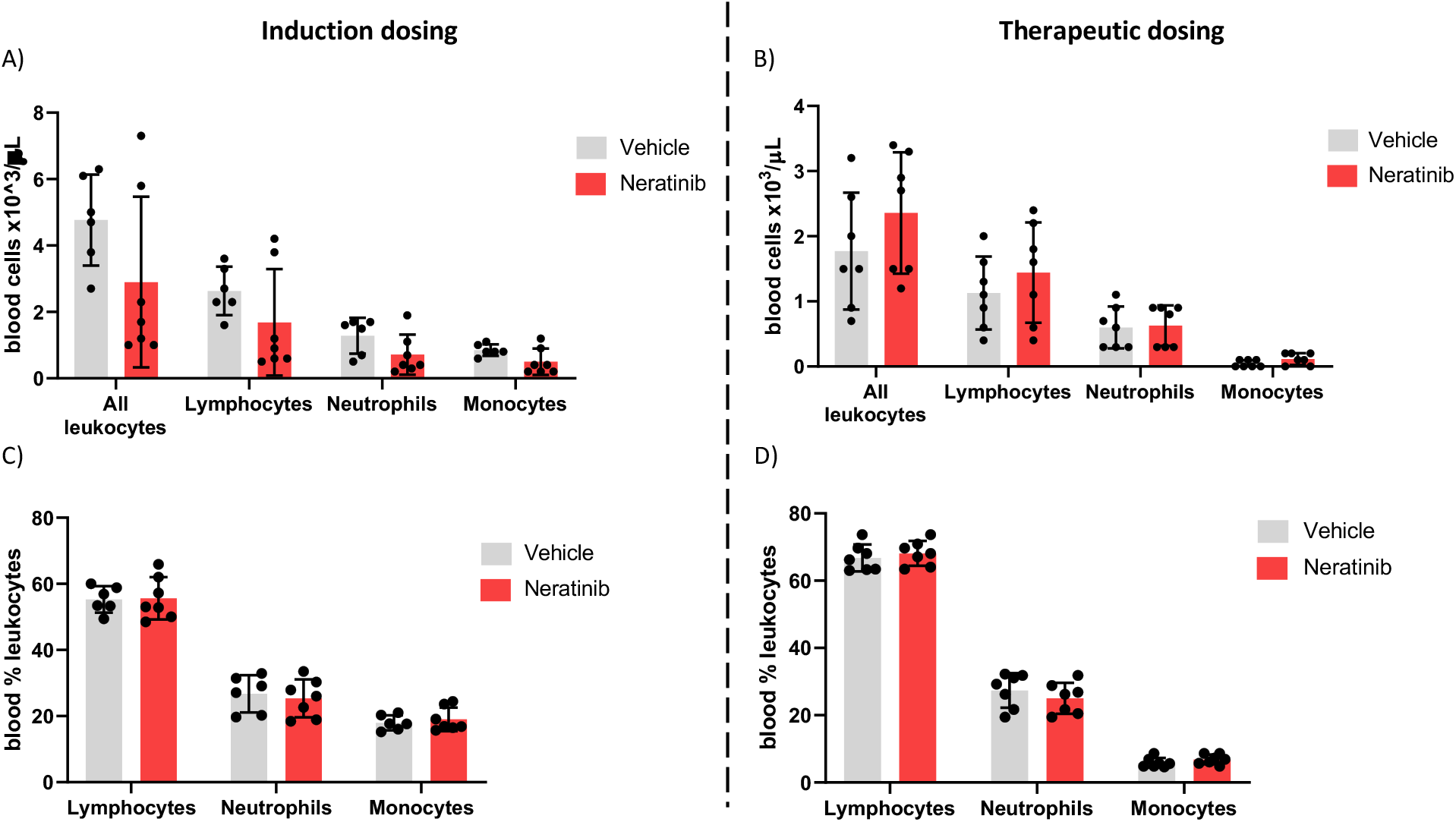
Neratinib does not alter the numbers or distribution of circulating leukocytes in chronic mouse models of lung disease. Leukocytes in mouse whole blood samples were analysed using an automated haematology blood analyser, which counts total leukocytes, as well as differentiating between lymphocytes, neutrophils and monocytes; the percentages of each leukocyte subset are also analysed. No differences in the number of leukocytes, or their subsets, were identified between vehicle and neratinib treated groups in the induction dosing (A) or therapeutic dosing (B) model, and similarly no differences in the percentages of lymphocytes, neutrophils and monocytes were found between treatment groups in the induction (C) or therapeutic (D) dosing model. Each data point represents data from one mouse; bars show mean ± standard deviation. Unpaired t tests used to compare treatment groups within each cell type, p values indicated where appropriate.

## Discussion

The suppression of neutrophil apoptosis and the release of neutrophilic inflammatory mediators in injured tissue are essential for efficient pathogen clearance, and individuals with defective neutrophils or chronic neutropenia experience life-threatening infections (Dinauer, 2020; Donadieu et al., 2011). However in chronic inflammatory diseases such as COPD, the continual release of neutrophilic reactive oxygen species, elastase and other proteolytic enzymes destroys alveolar epithelial cells, drives emphysematous changes, and damages the alveolar attachments that support bronchioles, resulting in their premature collapse (Barnes et al., 2015; Craig et al., 2017). Modifying neutrophils in this disease therefore may have therapeutic potential, although currently there are no therapeutics that do this. A number of studies have shown the beneficial effects of inducing apoptosis and reducing the functional activity of neutrophils, as well as increasing the rate of efferocytosis by macrophages, in various experimental inflammatory models (Chello et al., 2007; Ren et al., 2008; Rossi et al., 2006). Here we show that neratinib, an ErbB inhibitor used to treat breast cancer, may be able to reduce acute inflammation in the lungs, by reducing the number of neutrophils corpses, and increasing rates of efferocytosis.

The rapid removal of apoptotic neutrophils from an inflammatory environment before they become necrotic is crucial to prevent the highly histotoxic intracellular contents inducing tissue damage and further inflammation (Vlahos and Bozinovski, 2014). The process of efferocytosis itself is also a key step in inflammation resolution, skewing macrophages toward an anti-inflammatory phenotype in which they release IL-10, TGFβ and prostaglandin E2 to facilitate wound healing, and supress the production of pro-inflammatory cytokines such as IL-1β, CXCL-8, TNFα, and GM-CSF (Fadok et al., 1998; Sun et al., 2015; Wynn and Vannella, 2016). Interestingly, research using cell therapy (injected stem cells) to ameliorate inflammatory disease in mouse models has suggested that apoptosis of the injected cells, and subsequent efferocytosis by macrophages, is causing the immunosuppressive effects, rather than bioactivity of the stem cells themselves (Pang et al., 2021). In mouse models of pneumococcal pneumonia, instilled apoptotic macrophages are engulfed by resident tissue macrophages, which resulted in decreased neutrophilic inflammation and inflammatory cytokine levels in the lungs, and lower probabilities of the mice becoming bacteremic (Marriott et al., 2006).

It is unclear in our studies whether neratinib is directly regulating macrophage efferocytosis, or upregulating apoptosis and thereby increasing the amount of material available to engulf. There is very little literature suggesting that ErbB signalling has a role in efferocytosis, although a recent study in *Drosophila* from our group shows that overexpression of the cardinal EGF ligand, Spitz, impairs macrophage-mediated apoptotic cell clearance (Tardy et al., 2021). While this supports our current findings, there are likely to be a number of mechanisms at play, including EGF acting as a chemoattractant to “distract” the macrophages from apoptotic cell clearance, or EGFR signalling from other tissues polarising the macrophages towards a phenotype less efficient at efferocytosis (Tardy et al., 2021).

The binding of macrophage receptors to apoptotic markers such as phosphatidylserine activates intracellular macrophage signalling pathways including PI3K-AKT, STAT-SOCS, and Rho GTPases, and supresses NFκB and IFNα/β signalling, resulting in the engulfment of the apoptotic cell (Elliott et al., 2017). Our phosphoproteomic analysis using Reactome identified an enrichment in the Rho GTPase cycle, a known controller of efferocytosis and phagocytosis (Kim et al., 2017; Kinchen and Ravichandran, 2007). Rho GTPases are regulated by GTPase-activating proteins (GAPs) and guanine nucleotide exchange factors (GEFs) (van Buul et al., 2014). Our phosphoproteomic analysis of human neutrophils identified three of these phosphorylated proteins in the neratinib-enriched dataset (ARHGAP1, ARHGEF2 and MYO9B) and one in the DMSO-enriched dataset (ARHGAP18). RhoA, which inhibits the Rho GTPase cycle and thus efferocytosis, is itself inhibited by the cholesterol-lowering drugs statins (Rikitake and Liao, 2005). Treatment with lovastatin upregulated efferocytosis in mouse lungs *in vivo*, and also in human macrophages from patients with COPD *in vitro* (Morimoto et al., 2006). As we carried out the phosphoproteomics experiments with human neutrophils, it is not possible to tell if neratinib is regulating these pathways in macrophages in our acute lung injury mouse model, and further work would be required to draw conclusions.

Rho GTPases regulate other neutrophil functions including adhesion, chemotaxis and recruitment (McCormick et al., 2019). Our phosphoproteomic analysis indicated neratinib regulates actin filament assembly and cytoskeletal organisation, which also control migration. In the zebrafish larvae, neratinib treatment resulted in a reduction in neutrophil numbers at the tail fin injury site, suggesting an impairment in neutrophil migration. ErbBs are known to regulate migration, with research demonstrating that overamplification of ErbB receptors on tumour cells can induce epithelial-mesenchymal transition (leading to metastasis), migration, and tumour invasion (Appert-Collin et al., 2015). Neratinib specifically reduces the migration of gastric adenocarcinoma cells *in vitro* (Hamzehlou et al., 2019). Our data suggest that neratinib-induced inhibition of migration might be due to the regulation of Rho GTPase pathways, although again further validation is required to conclude this.

The mechanism by which neratinib is promoting apoptosis of neutrophils *in vitro* is still unclear, although the phosphoproteomic analysis has elucidated several potential candidates. Nucleophosmin 1 (NPM1) is a protein with a range of intracellular functions including ribosome biogenesis, protein chaperoning, histone assembly and regulation of the tumour suppressor p53, and is often mutated or overexpressed in cancer cells, contributing to carcinogenesis (Grisendi et al., 2006). Our data show that a pharmacological inhibitor of NPM1 induces neutrophil apoptosis, both alone and in the presence of neratinib, suggesting these two inhibitors may be inducing apoptosis in neutrophils via the same mechanism. We detected phosphorylation of nucleophosmin 1 (NPM1) at serine 4 and serine 10 in 3/4 DMSO treated samples and of 0/5 neratinib treated samples, suggesting a potential role in the induction of apoptosis with neratinib treatment. Other sources have linked these specific phosphorylation events to cell death, for example irradiated basal epithelial cells show dephosphorylation of NPM1 at serine 4 (Wiesmann et al., 2019). Inactivation of the serine 10 and serine 70 phosphorylation sites of NPM1 induces cell cycle arrest in mutant lymphoblastoid cells, and also positively regulates the activity of cyclin-dependant kinase 1, a key regulator of the cell cycle (Du et al., 2010).

Targeting EGFR signalling as a therapeutic strategy for inflammatory lung diseases has been investigated by others, although not in the context of modifying neutrophils specifically. Bronchial epithelial cells from patients with COPD have increased production of CXCL8 and increased phosphorylation of EGFR and AKT, however all were reduced *in vitro* with erlotinib, a clinical EGFR inhibitor (Ganesan et al., 2013). The binding of the EGFR ligand amphiregulin was shown to be essential for TGFβ-dependant pulmonary fibrosis in mouse models (Andrianifahanana et al., 2013). The EGFR signalling pathway was also shown to contribute to the loss of muscle function that many patients with COPD experience (Ciano et al., 2019). Rhinovirus infection upregulates mucin production in the airways via the NFκB and EGFR pathways, and this was supressed in mouse models of COPD exacerbations using an EGFR inhibitor (Hewson et al., 2010; Singanayagam et al., 2022). A clinical trial of an inhaled EGFR antagonist however did not reduce mucus production in patients with COPD (Woodruff et al., 2010). Although EGFR inhibitors have to date only been approved for cancer treatment, other research does suggest they have potential benefit for inflammatory lung diseases such as COPD.

The results from our chronic mouse models of lung inflammation suggest that neratinib (as dosed here) is less effective over longer time periods. The reduction of cytokines KC and IL-6 in the induction dosing model of neratinib treatment is promising, although it is unclear if this parameter alone would confer therapeutic benefit. The reduction in KC is particularly interesting as this cytokine is a key recruiter of neutrophils to sites of inflammation or infection (Sawant et al., 2016). Other murine acute lung injury models show that depletion of KC correlates with decreased early neutrophil recruitment and improved histopathology scores, suggesting a potential therapeutic benefit (Dunn et al., 2018). Blood biomarkers of cytokines such as IL-6 have also been shown to correlate with disease severity in patients with COPD (Bradford et al., 2017). We measured no change in the number or proportion of neutrophils in BAL, and it may be that other inducers of neutrophil recruitment in mice such as LTB4 and CXCL5 (Grespan et al., 2008; Saiwai et al., 2010) render the abrogation of KC redundant. We also measured no changes in cytokine levels with the therapeutic dosing protocol of neratinib, suggesting that treatment is required earlier in the course of disease to show efficacy.

Since neratinib showed the most efficacy in the acute lung inflammation mouse model, it may be that for COPD patients this drug would most benefit those experiencing exacerbations, rather than as a long-term treatment option. Further pre-clinical models and clinical trials would ultimately be required to determine any potential benefit; however, this work has given additional evidence to support neratinib repurposing, narrowed down the inflammatory context in which it may have therapeutic potential, and further defined the intracellular regulators of by which neratinib exerts its anti-inflammatory effects.

## Materials and methods

### Zebrafish husbandry and treatment with pharmacological agents

All zebrafish were raised and maintained according to standard protocols (Nüsslein-Volhard and Dahm, 2002) in UK Home Office approved aquaria at the University of Sheffield, according to institutional guidelines. The transgenic zebrafish line *TgBAC(mpx:EGFP)i114*, in which neutrophils express green-fluorescent protein (GFP) under the myeloid-specific peroxidase (*mpx*) promoter, was used for all experiments (Mathias et al., 2006; Renshaw et al., 2006). For the treatment of zebrafish larvae with neratinib, larvae at 2 days post-fertilisation (dpf) were immersed E3 media containing dissolved neratinib HKI-727 (Selleck, Houston, Texas, S2150), or equivalent volume of DMSO as a control, and incubated at 28°C. This was carried out in 6-well plates, each well containing 6mL of E3 and a maximum of 15 larvae. For analysis, each larva counted as one biological replicate. Experiments were repeated three times with different batches of larvae born from different tanks of adult zebrafish.

### Whole body neutrophil counts in zebrafish larvae

To enumerate the total number of neutrophils across the whole body of zebrafish larvae, larvae were treated for 16 hours (overnight) with neratinib or DMSO, anaesthetised with tricaine, then mounted in low melting-point agarose for imaging. Images were acquired using a Nikon Eclipse TE2000 U (Nikon, Tokyo, Japan) inverted compound fluorescence microscope with NIS-Elements software. Neutrophils were enumerated manually by GFP expression.

### Tail fin transection model of injury-induced inflammation in zebrafish larvae

To assess neutrophil number at the site of inflammation, a tail fin transection model of injury-induced inflammation was used. After 16 hours of treatment with neratinib or DMSO, larvae were anaesthetised using tricaine, and complete transection of the tail fin was carried out using a sterile scalpel as described previously (Lieschke et al., 2001; Renshaw et al., 2006), at a site distally adjacent to the circulatory loop. After transection, larvae were transferred to fresh E3 containing neratinib or DMSO. At 4 and 8 hours post injury, larvae were anaesthetised with tricaine and neutrophils at the injury site enumerated based on GFP expression, using either a Leica Fluorescence Stereo Dissecting Microscope (Leica, Wetzlar, Germany) or a Nikon Eclipse TE2000 U with NIS-Elements software.

### Isolation of neutrophils from human whole blood

*In vitro* experiments were conducted on neutrophils isolated from peripheral blood samples of healthy donors, in compliance with the guidelines of the South Sheffield Research Ethics Committee (reference number STH13927). The volume of blood required was calculated prior to collection according to experimental requirements, based on approximately 2 million isolated neutrophils per 1 mL blood. Peripheral blood samples were obtained by venepuncture and collected into a 50mL syringe, then immediately transferred into a 50mL Falcon tube containing EDTA, 122μL per 10mL blood.

Isolation of neutrophils was carried out EasySep™ Direct Human Neutrophil Isolation Kit (Stem Cell Technologies, Vancouver, Canada, 19666), an immunomagnetic negative selection kit, as per the kit instructions. In brief, the antibody Isolation Cocktail and magnetic RapidSpheres (25μL each per 1mL blood) were added to the blood sample and incubated for 5 minutes, and the tube then placed in the EasySep magnet for 5 minutes. After incubation, the enriched neutrophil sample was removed and transferred to a fresh tube, leaving non-neutrophils bound to the antibody/bead complex immobilised against the side of the original tube by the magnet. Two more magnet incubations of the enriched neutrophil sample were carried out to further purify the neutrophils. The final purified neutrophil sample was washed by centrifugation at 300g for 6 minutes, and resuspended in 10mL of phosphate-buffered saline (PBS). A haemocytometer counting chamber was used to conduct cell counts, and the washed neutrophils resuspended in RPMI 1640 media (Lonza, Basel, Switzerland, BE12-702F), supplemented with 10% fetal bovine serum (FBS) (Gibco, Waltham, Massachusetts, 10500-064) at a concentration of 5×10^6^ cells/mL.

### Treatment of human neutrophils with pharmacological inhibitors and assessment of apoptosis

Isolated human neutrophils were added to a 96-well flexible untreated polyvinyl chloride general assay plates (Corning, New York, CLS2592), which minimise neutrophil adherence and activation. Plates were prepared in advance of the addition of neutrophils, with each well containing 50μL neratinib HKI-272 (Selleck, S2150), EX527 (SIRT1 inhibitor - Santa Cruz Biotechnology, Dallas, Texas), NSC348884 (NPM1 inhibitor - Selleck), or DMSO at 2X desired concentration in RPMI 1640 + 10% FBS media. Two technical replicates (wells) of each drug and concentration were generated. Immediately after isolation, 50μL neutrophils at a concentration of 5×10^6^/mL were added to each well, diluting the pharmacological agent to the desired concentration. Plates were incubated at 37°C in 5% CO_2_ for 6 hours.

For the fixation, staining and imaging of neutrophils, cytocentrifugation was used to transfer a monolayer of cells onto glass microscopy slides, referred to as “cytospin slides”. Cells in each well were transferred to a cytospin funnel secured with a clamp onto a glass microscope slide, and centrifuged in a Shandon Cytospin 4 cytocentrifuge (Thermo Electron Corporation, Waltham, Massachusetts) at 300rpm for 3 minutes. Neutrophils (now immobilised on the microscope slide) were fixed with a drop of 100% methanol, and stained by submersion in Kwik-Diff Solution 2 (Thermo Electron Corporation, 9990706) followed by Kwik-Diff Solution 3 (Thermo Electron Corporation, 9990707). Excess stain was washed from the slide using tap water, and once dry a drop of DPX (Sigma Aldrich, St Louis, Missouri, 44581) was added directly onto the cells, and a coverslip placed carefully on top. Slides were left in the fume hood for at least 24 hours to allow the DPX to set, then imaged using a Nikon Eclipse TE300 inverted light microscope with 100X magnification oil immersion lens. Apoptotic neutrophils were identified based on nuclear morphology, which in apoptotic neutrophils is rounded and condensed, in comparison to the distinctive multi-lobed nuclei of healthy neutrophils. To calculate the percentage apoptosis, 300 neutrophils per cytospin slide were counted and recorded as healthy or apoptotic. As two technical replicates were generated per condition, a mean percentage apoptosis was calculated to generate one datapoint per condition per blood donor.

### Mass spectrometry-based phosphoproteomics analysis of human neutrophils treated with neratinib

Neutrophils isolated from blood of 5 healthy volunteers, as described above. Neutrophils from each donor were split into two samples of 15 million cells each and treated with 25μM neratinib (Sellek) or equivalent concentration (v/v) DMSO, and incubated for 1 hour at 37°C and 5% CO_2_. All samples were then treated with 500μM db-cAMP (Sigma) for 30 minutes in the same incubation conditions. After incubation, samples were centrifuged at 400g for 3 minutes at 4°C and the cell pellet resuspended in 1mL ice-cold PBS. Cells were centrifuged again at 400g for 3 minutes at 4°C, the PBS supernatant removed, and cell pellets stored immediately at −80°C.

To extract protein from samples, the following lysis buffer was added to the neutrophil pellets: 5% SDS, 50mM TEAB buffer, 50mM sodium fluoride, 50mM β-glyercophosphate, 10mM sodium orthovanadate, 1mM PMSF, and 5% Protease Inhibitor Cocktail Set III (Calbiochem, San Diego, California, 535140) made up in HPLC-grade water. DNA was sheared from the lysed cells using a homogeniser. Samples were incubated at 70°C for 15 minutes, homogenised again and incubated for a further 5 minutes, or until no cellular material was visible. Samples were centrifuged at 16,000g to pellet any cell debris, and protein supernatants transferred to fresh Eppendorf tubes. Samples were reduced with 10mM TCEP and alkylated with iodoacetamide.

Protein purification was carried out using suspension trapping (S-Trap™) columns which contain a protein-binding matrix (Protifi, Long Island, New York, K02-Micro-10). To each 120μL protein extract, 12μL 12% phosphoric acid and 840μL binding buffer (90% methanol, 100mM TEAB buffer) was added. Samples were transferred to the top chamber of an S-Trap™ spin column and centrifuged at 4000g for 15 seconds. The proteins (now “trapped” in the matrix) were washed four times with 400μL binding buffer and centrifuging as before to elute any impurities. Proteins were digested into peptides with 12μL of 1:10 Trypsin Gold (Promega, Madison, Wisconsin, V528A) in 50mM TEAB per sample, and incubated at 47°C for 1 hour. Peptides were then eluted from the S-Trap™ matrix with 80μL 50mM TEAB, centrifuging at 1000g for 30 seconds, followed by 80μL 0.2% formic acid solution and centrifuging again at 1000g for 30 seconds. Peptides were desalted using Sep-Pak^®^ Light C18 Cartridge (Waters, Wilmslow, UK, WAT023501) and dried down using a SpeedVac (Thermo Scientific).

Immobilised metal affinity chromatography (IMAC) was used to enrich for phosphorylated peptides, using MagReSyn^®^ Ti-IMAC beads (ReSyn Biosciences, Gauteng, South Africa, MR-THP002). Peptides were firstly resuspended in IMAC loading buffer (1M glycolic acid, 80% acetonitrile, 5% trifluoroacetic acid) and centrifuging at 1000rpm for 5 minutes. Beads were placed on a magnetic rack and washed with IMAC loading buffer, after which peptide samples were added to the beads and incubated for 20 minutes. After centrifugation at 1000 rpm, the supernatant was removed and three washes were carried out using 100μL IMAC loading buffer per wash. Enriched phosphorylated peptides were eluted from the beads with 80μL 1% ammonia, then acidified with 40μL 10% trifluoroacetic acid.

Phosphopeptides were analysed by high performance liquid chromatography-mass spectrometry, using HPLC column Acclaim^®^ PepMap 100 C18 nano/capillary BioLC (ThermoFisher Scientific, 164535), EASY-Spray column (ThermoFisher Scientific, ES803), and analysis on an Orbitrap Elite™ Hybrid Ion Trap. Raw data was analysed using MaxQuant version 1.6.10.43 software. Peptide spectra were searched against a human UniProt fasta file (downloaded May 2019) using the following search parameters: digestion set to Trypsin/P with a maximum of 2 missed cleavages, oxidation (M), N-terminal protein acetylation and phosphorylation (STY) as variable modifications, cysteine carbamidomethylation as a fixed modification, match between runs enabled with a match time window of 0.7 min and a 20-min alignment time window, label-free quantification was enabled with a minimum ratio count of 2, minimum number of neighbours of 3 and an average number of neighbours of 6. A first search precursor tolerance of 20 ppm and a main search precursor tolerance of 4.5 ppm was used for FTMS scans and a 0.5 Da tolerance for ITMS scans. A protein FDR of 0.01 and a peptide FDR of 0.01 were used for identification level cut-offs and an FLR of 5% for phosphosite localisation. Statistical analysis of the phosphorylation site data was performed using Perseus version 1.6.10.50. Phosphorylation site intensities were transformed by log2(x), normalised by subtraction of the median value, and individual intensity columns were grouped by experiment. Phosphorylation sites were filtered to keep only those with a minimum of 3 valid values in at least one group. The distribution of intensities was checked to ensure standard distribution for each replicate. Missing values were randomly imputed with a width of 0.3 and downshift of 1.8 from the standard deviation. To identify significant differences between groups, two-sided Student’s t-tests were performed with a permutation-based FDR of 0.05.

### Mouse husbandry and models of lung inflammation

Approval for working with mammalian models was authorised by the Home Office under the Animals (Scientific Procedures) Act 1986 under project license P4802B8AC held by Dr Helen Marriott, and personal licenses (PIL) held by Carl Wright, Sam McCaughran and Dr Helen Marriot, and reviewed by the Animal Welfare and Ethical Review Body at the University of Sheffield. All housing parameters conformed to the Code of Practice for the housing and care of animals bred, supplied or used for scientific procedures. Food (Teklad 2018, Envigo, Indianopolis, Indiana) and water were given ad-lib, and animals kept on a 12h light-dark cycle. To adhere to humane end-points, if mice appeared in distress and were not comfortably breathing 24 hours after a procedure, or if 20% weight loss were reached, mice were culled to prevent excessive suffering. Female C57BL/6J mice, aged 9-12 weeks and weighing 16-20g at the start of each study were used for all experiments.

In the acute lung injury study, 12 mice were anaesthetised with gaseous isoflurane and administered 7μg lipopolysaccharides (LPS) from *E. Coli* O26:B6 (LPS – Sigma, L8274) in 50μL PBS. Immediately after, 6 mice were administered 200μL vehicle (0.5% methylcellulose + 0.4% Tween-80 + 1% DMSO) by oral gavage, and the remaining 6 mice administered 20mg/kg neratinib (ApexBio Technology, Houston, Texas, A8322) dissolved in vehicle, by oral gavage. Mice were given immediate heat support and observed regularly by experienced PIL holders. After 48 hours, all mice were sacrificed by terminal anaesthesia, by intraperitoneal administration of 100μL pentobarbitone (100mg/mL), and subject to bronchoalveolar lavage (3x 1mL administrations of PBS).

In the two chronic lung injury studies, 16 mice per study were administered 7μg LPS (as above) combined with 1.2U porcine pancreatic elastase (Merck, Darmstadt, Germany, 324682) in 50μL PBS, by intranasal delivery as above. This was carried out weekly for 4 weeks, i.e. on days 0, 7, 14, 21. For the induction dosing study, neratinib or vehicle were administered to the mice as above (8 mice per treatment group), immediately after each LPS/elastase administration. For the therapeutic dosing model, neratinib or vehicle doses were given on days 14, 15, 16, 17, 18, 21, 22, 23, 24, 25. In both chronic disease models, all mice were sacrificed by terminal anaesthesia on day 28, bronchoalveolar lavage fluid was collected along with blood samples by inferior vena cava bleed.

### Preparation of mouse bronchoalveolar lavage samples for cell counting, microscopy and ELISA

Cells in bronchoalveolar lavage (BAL) samples were placed immediately on ice after collection. A haemocytometer counting chamber was utilised to calculate the total number of cells in each sample. In a centrifuge pre-cooled to 4°C, BAL samples were centrifuged at 400g for 5 minutes to pellet the cells, and supernatant removed and stored immediately at −80°C for later analysis by ELISA. The remaining cells were resuspended in ice-cold PBS at a concentration of 2 million/mL. Cytospin funnels were assembled, and 50μL of each BAL sample transferred to each funnel, to which 50uL FBS was added to prevent the cells from breaking during centrifugation. A cytocentrifuge was used to transfer the cells to microscope slides, and slides were fixed and stained with Kwik-Diff as described above for human neutrophils.

### Preparation of mouse bronchoalveolar lavage for flow cytometry

The cells remaining in each BAL sample were prepared for flow cytometry. For all mouse studies, cells were stained with FITC anti-mouse Ly6G/Ly6C antibody (Biolegend, San Diego, California, 108405) to detect neutrophils; PE-Annexin V (Biolegend, 640908) which binds to apoptotic cells; and TO-PRO™-3 Iodide (Invitrogen, Waltham, Massachusetts, T3605) a vital dye that only binds to dead cells in which the plasma membrane is broken. All samples were stained with all three markers, and an additional four samples were generated from the BAL samples with the most cells, for one unstained control and three single-stain controls. All samples were centrifuged at 400g for 3 minutes at 4°C to pellet the cells, and resuspended in 50uL FITC-Ly6G staining solution diluted 1:200 in FACS buffer (PBS + 10% FBS), or 50uL FACS buffer for controls. Samples were incubated on ice, in the dark for 20 minutes. Samples were centrifuged at 400g for 3 minutes at room temperature and resuspended 50μL in PE-Annexin V antibody diluted 1:20 with Annexin V Binding Buffer (BioLegend, 422201), or 50uL Annexin V Binding Buffer only for controls. Samples were incubated in the dark at room temperature for 20 minutes, after which 5uL of TO-PRO-3 solution (diluted 1:1000 in Annexin Binding Buffer) was added to each sample. Samples kept in the dark at room temperature while the flow cytometry analyser was set up (approximately 10 minutes). When samples were ready for analysis, 300uL of Annexin Binding Buffer was added to each sample to ensure an adequate volume for analysis. Samples were run on a BD LSRII Flow Cytometer (BD Biosciences, Franklin Lakes, New Jersey) using BD FACSDiva™ software, and data analysis carried out using FlowJo software (BD Biosciences).

### Analysis of mouse blood samples using an automated haematology analyser and preparation of plasma for ELISA

Blood was collected into EDTA-coated tubes to prevent clotting, and mixed gently. The majority of each sample was transferred to a fresh labelled Eppendorf tube, leaving 50μL blood from in each EDTA tube for analysis by an automated haematology blood analyser. Blood samples from the first chronic lung disease mouse study (induction neratinib dosing) were analysed using a Sysmex KX-21N™ (Hyogo, Japan), that had been adapted in-house to measure parameters for mouse blood cells, including leukocyte concentration, and differential detection of neutrophils, monocytes and lymphocytes. For the second chronic lung disease study (therapeutic neratinib dosing), blood samples were analysed using a scil Vet abc Plus^+^ (Scil, Viernheim, Germany) automated haematology analyser, which is programmed to analyse blood samples from a range of animals including mice.

The remaining blood samples were centrifuged at 350g for 10 minutes at 4°C to separate the plasma from the blood cells. The upper plasma layer was transferred to a fresh Eppendorf and stored immediately at −80°C, for later analysis by ELISA.

### IL-6 and CXCL1/KC ELISA

An enzyme-linked immunosorbent assay (ELISA) was used to determine the concentration of cytokines in bronchoalveolar lavage fluid and plasma samples from mice. ELISA kits used were Mouse IL-6 DuoSet ELISA (R&D Systems, Minneapolis, Minnesota, DY406-05) and Mouse CXCL1/KC DuoSet ELISA (R&D Systems, DY453-05), and the assays carried out as per the kit instructions. All BAL samples were run neat. Two technical replicates of all samples and standards were used. Plates were analysed on a Thermo Scientific Varioskan^®^ Flash microplate reader at 450nm, with wavelength correction at 540nm. Cytokine concentration was calculated using interpolation of a 4-parameter logistic sigmoidal standard curve, as suggested by the kit instructions.

### Statistical analysis of data from mouse studies

Statistical analysis on the data collected from mouse studies compared the vehicle and neratinib treatment groups. Datasets contained a single data point for each mouse, which in some cases was calculated as an average of technical replicates within the assay (e.g. ELISA), but others were from a single measurement (e.g. leukocyte number in blood samples). An unpaired t test was used to compare neratinib vs vehicle treatment groups.

## Acknowledgements

We thank the Biological Services Aquarium staff and the Biological Services Unit staff for their assistance with zebrafish and mouse husbandry. We would also like to thank the following facilities at the University of Sheffield: the Wolfson Light Microscopy Facility, the Medical School Flow Cytometry Core Facility, and the biOMICS Mass Spectrometry Facility. Lastly, we would like to thank the volunteers who donated blood for this research.

This work was supported by a Medical Research Council (MRC) Programme Grant to S.A.R (MR/M004864/1), a Rosetrees Trust grant to L.R.P (100179), and the Faculty of Medicine, Dentistry and Health Doctoral Academy Scholarship to K. D. H.

## Figure Supplements

**Figure 2 - figure supplement 1.**
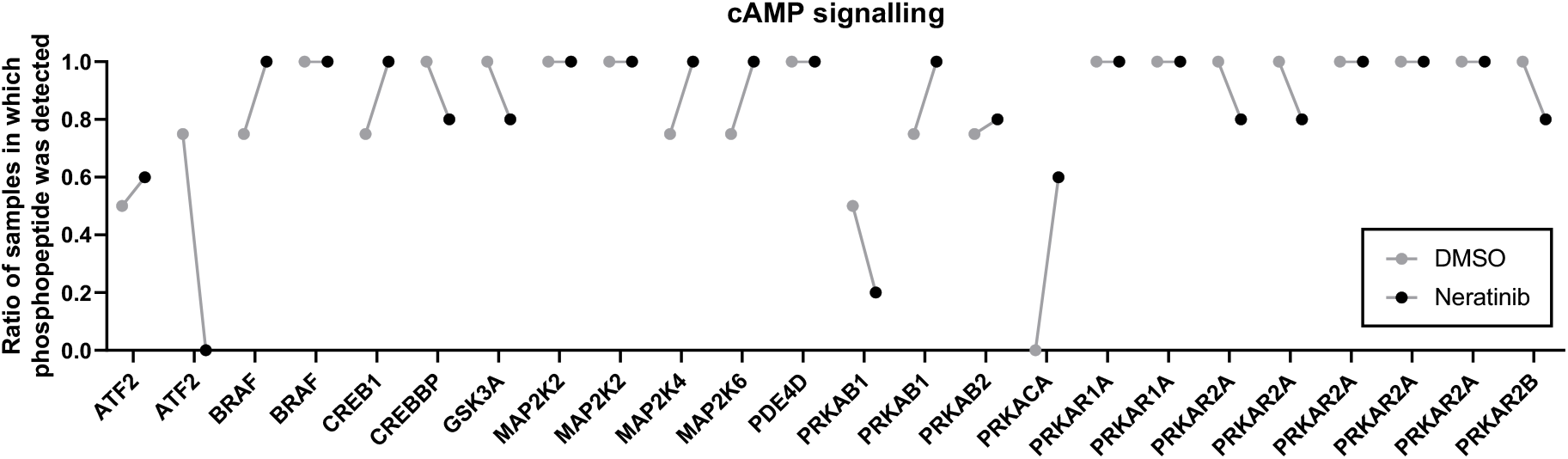
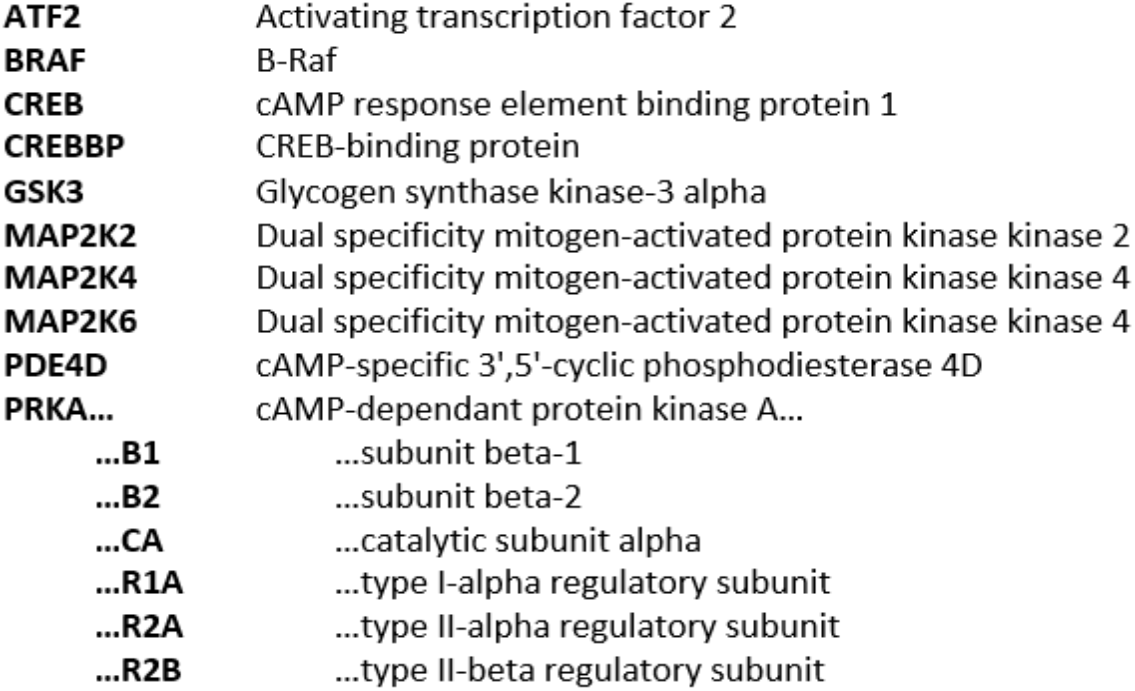
Validation of the dataset by identifying phosphorylated proteins downstream of cAMP signalling pathways. Phosphorylated proteins downstream of cAMP signalling pathways were identified from the dataset. cAMP binds to the regulatory subunit of inactive protein kinase A (PKA), resulting in the dissociation and activation of catalytic PKA subunits, which in turn phosphorylate a number of downstream targets, such as BRAF, GSK3A and MAPK proteins, and transcription factors AFT and CREB in the nucleus. cAMP is also converted to 5’ AMP by PDE enzymes. The data was analysed by noting the number of samples in which a phosphorylated peptide mapping to a protein was detected in the DMSO and neratinib treatment groups. This is represented as a ratio of the total number of samples in each treatment group. For example, a phosphorylated peptide mapping to CREB1 was detected in 3/4 (0.75) DMSO treated samples and 5/5 (1) neratinib treated samples. In some cases, multiple phosphorylated peptides mapping to the same protein were identified.

**Figure 2 - figure supplement 2.**
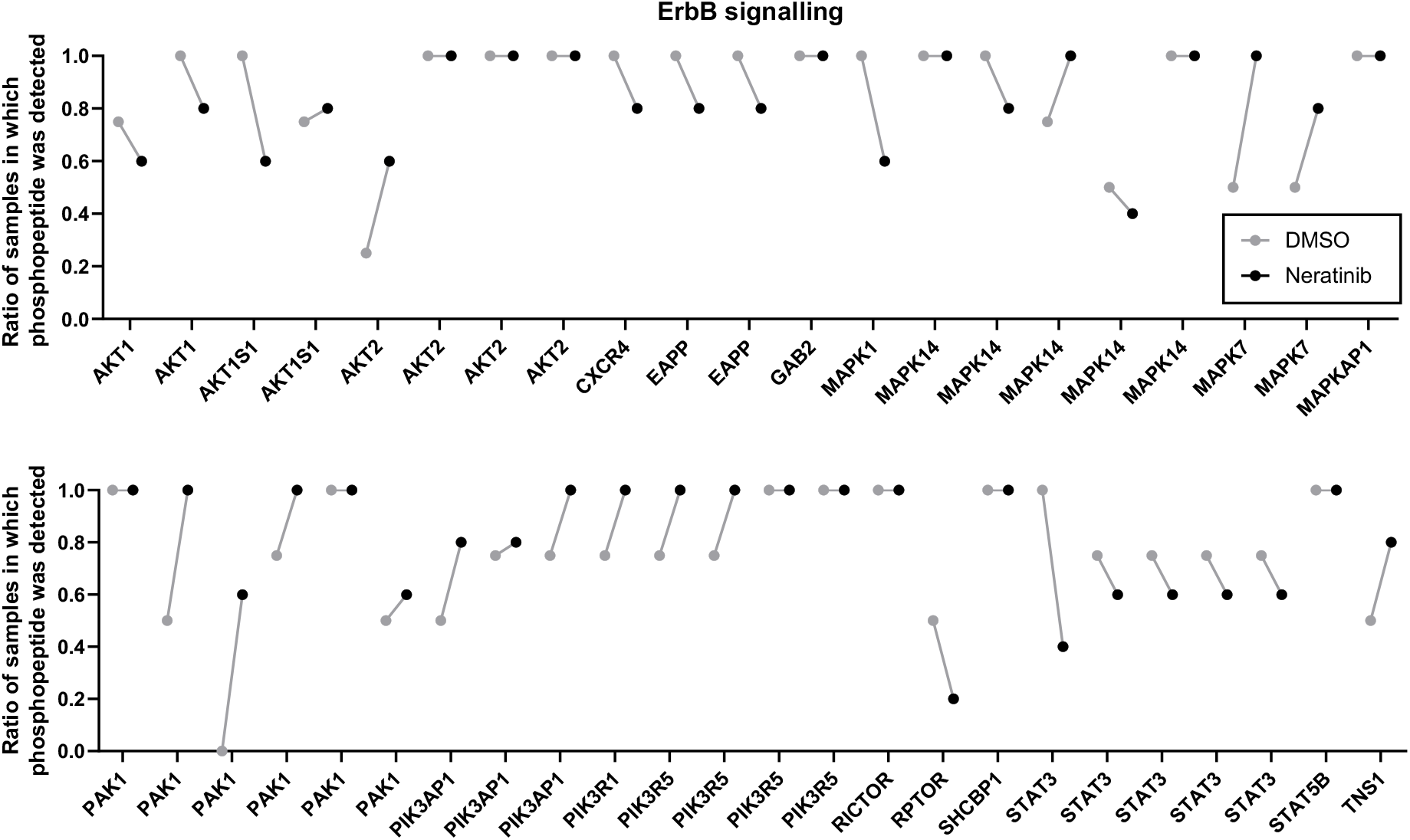
Detection of phosphorylated proteins downstream of ErbB signalling pathways. Phosphopeptides mapping to proteins downstream of ErbB signalling pathways were identified within the dataset and shown as a ratio of the number of samples in each treatment group it was detected in, out of 4 DMSO treated samples and 5 neratinib treated samples. In some cases, multiple phosphorylated peptides mapping to the same protein were identified.

**Figure 2 - figure supplement 3.**
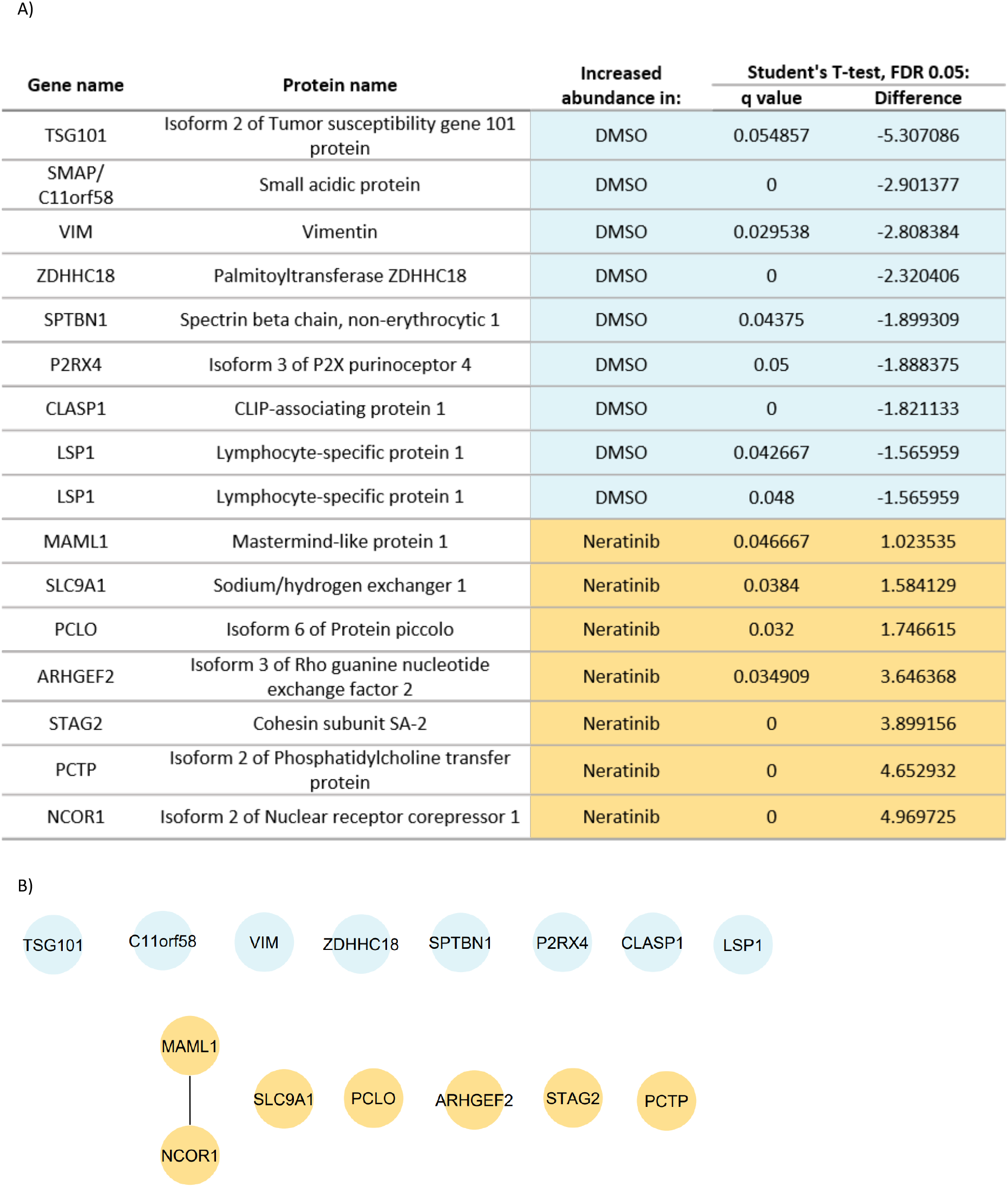
Statistical analysis of phosphopeptide abundance between treatment groups. Statistical analysis of the phosphoproteomics dataset identified 16 phosphopeptides with significantly different abundances between the neratinib and DMSO treated neutrophils, at 5% permutation-based false discover rate (FDR) using a paired student’s t test. This dataset was input into STRING, and one interaction between two phosphorylated proteins increased with neratinib treatment, MAML1 and NCOR1, was identified (B).

**Figure 2 - figure supplement 4.**
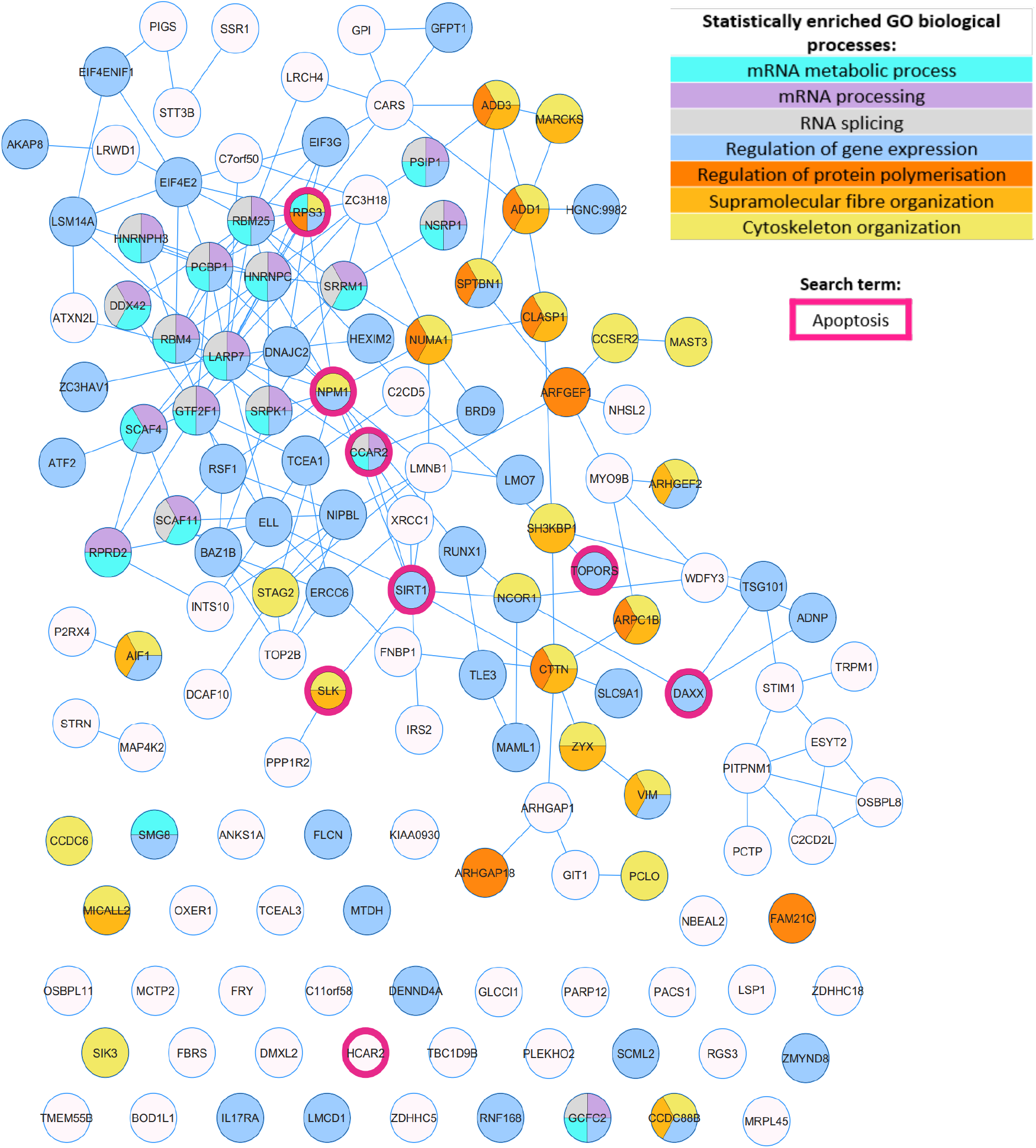
STRING analysis of phosphorylated proteins enriched and statistically regulated in both treatment groups. The DMSO- and neratinib-enriched phosphorylated proteins, and the statistically regulated phosphorylated proteins, were combined into one dataset and analysed in STRING. A selection of the biological processes that were identified as statistically enriched are highlighted in colour. Although not statistically enriched, a number of proteins are also involved in the regulation of apoptosis (pink outlines). Lines indicate interactions between proteins; some proteins have multiple interactions, whereas others have none.

**Figure 2 - figure supplement 5.**
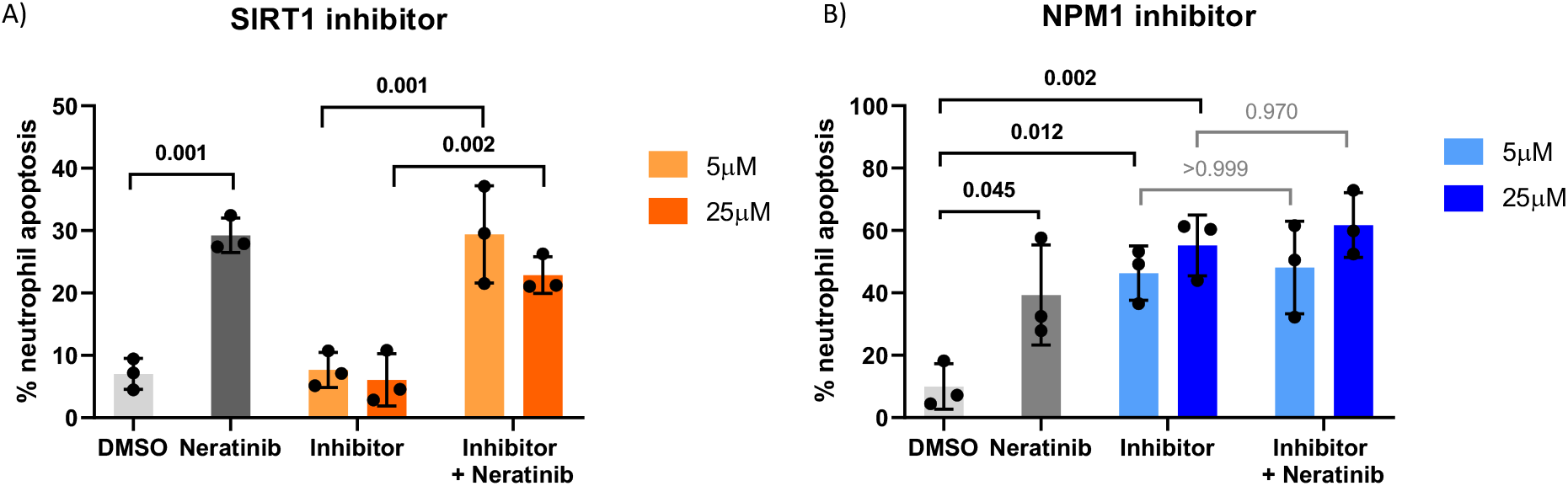
Pharmacological inhibition of candidates from the phosphoproteomics analysis reveal NPM1 as potentially being downstream of neratinib-induced neutrophil apoptosis. Human neutrophils were incubated with inhibitors of SIRT1 or NPM1, alone or in combination with 25 μM Neratinib, for 6 hours, after which levels of apoptosis were assessed by morphology. In both experiments, neratinib alone induced neutrophil apoptosis in comparison to the DMSO control, as expected. The inhibitor of SIRT1 did not induce human neutrophil apoptosis (A). The inhibitor of NPM1 did induce human neutrophil apoptosis (B), and no additional apoptosis was observed when this inhibitor was used in combination with neratinib. Each datapoint shows data from one healthy donor, bars represent mean ± standard deviation. One-way ANOVA with multiple comparisons used to calculate statistical significance, p values indicated.

**Figure 3 - figure supplement 1.**
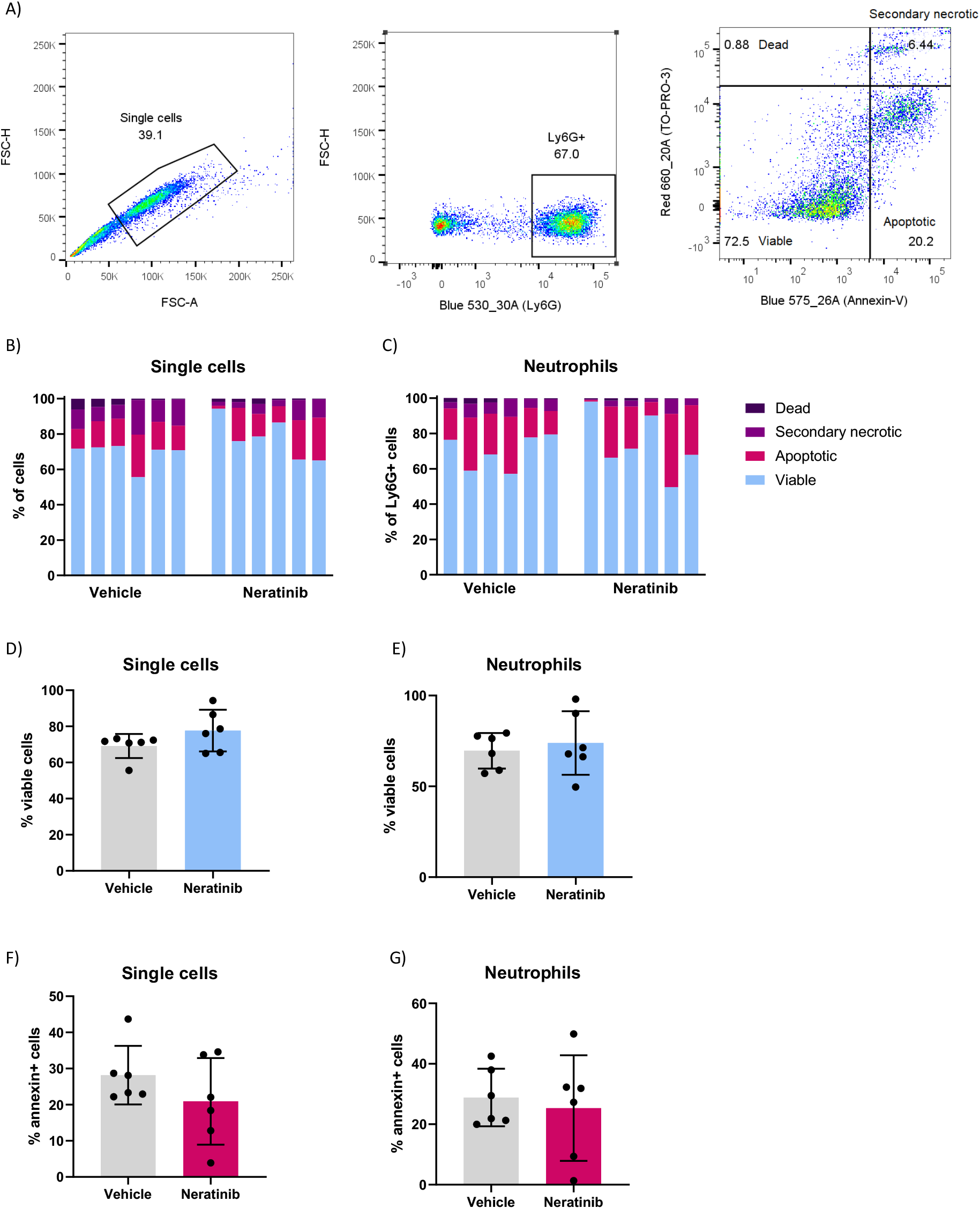
Flow cytometry analysis of BAL cells from the LPS-induced acute lung injury mouse model. Cells in BAL were analysed by flow cytometry. Single cells were initially gated (A, left panel), followed by Ly6G+ cells to identify neutrophils (A, middle panel). Annexin-V and TO-PRO-3 staining were then used to categorise cells as viable (negative for Annexin-V and TO-PRO-3), apoptotic (Annexin-V+), dead (TO-PRO-3+) and secondary necrotic (Annexin-V+ and TO-PRO-3+) (A, right panel). The single cells (B) and neutrophils (C) from each mouse BAL sample were categorised as such. Each bar represents data from one mouse. The percentage of viable cells (D) and viable neutrophils (E) was unchanged between treatment groups, as was the percentage of apoptotic cells (F) and apoptotic neutrophils (G). D-G: each data point represents data from one mouse; bars show mean ± standard deviation. Unpaired t tests used for statistical analysis; no significant differences found between vehicle and neratinib treatment groups.

Supplementary File 1. Phosphoproteomics dataset.

## Notes

### Competing Interest Statement

The authors have declared no competing interest.

